# Ldb1 and Rnf12-dependent regulation of Lhx2 controls the relative balance between neurogenesis and gliogenesis in retina

**DOI:** 10.1101/183285

**Authors:** Jimmy de Melo, Anand Venkataraman, Brian S. Clark, Cristina Zibetti, Seth Blackshaw

## Abstract

Precise control of the relative ratio of retinal neurons and glia generated during development is essential for visual function. We show that *Lhx2*, which encodes a LIM-homeodomain transcription factor essential for specification and differentiation of retinal Müller glia, also plays a critical role in the development of retinal neurons. Overexpression of *Lhx2*, and its transcriptional coactivator *Ldb1*, triggers cell cycle exit and inhibits both Notch signaling and retinal gliogenesis. *Lhx2/Ldb1* overexpression also induced the formation of wide-field amacrine cells (wfACs). In contrast *Rnf12*, which encodes a negative regulator of LDB1, is necessary for the initiation of retinal gliogenesis. We also show that LHX2 protein binds upstream of multiple neurogenic bHLH factors including *Ascl1* and *Neurog2*, which are necessary for suppression of gliogenesis and wfAC formation respectively, and activates their expression. Finally, we demonstrate that the relative level of the LHX2-LDB1 complex in the retina decreases in tandem with the onset of gliogenesis. These findings show that control of *Lhx2* function by *Ldb1* and *Rnf12* acts as a molecular mechanism underpinning the coordinated differentiation of neurons and Müller glia in postnatal retina.

**Significance Statement:** The molecular mechanisms that control the ratio neurons and glia that are generated by neuronal progenitors remain unclear. Here we show that Lhx2, a transcription factor essential for retinal gliogenesis, also controls development of retinal neurons. The Lhx2 coactivator Ldb1 promotes Lhx2-dependent neurogenesis, while the Lhx2 corepressor Rnf12 is necessary and sufficient for retinal gliogenesis. Furthermore, Lhx2 directly regulates expression of bHLH factors that promote neural development, which are necessary for Lhx2-dependent neurogenesis. Finally, we show that levels of the LHX2-LDB1 complex, which activates transcription, drop as gliogenesis begins. Dynamic regulation of Lhx2 activity by Ldb1 and Rnf12 thus controls the relative levels of retinal neurogenesis and gliogenesis, and may have similar functions elsewhere in the developing nervous system.

## Introduction

*Lhx2* is one of 12 genes that comprise the LIM class homeodomain (LIM-HD) family of transcription factors (TFs). *Lhx2* is dynamically expressed in multiple tissues, including discrete domains within the central nervous system (CNS) (1, 2). In the developing visual system, *Lhx2* activation is concurrent with patterning of the optic primordia and remains ubiquitous during formation of the optic vesicle and optic cup (1, 3). *Lhx2* is expressed in retinal progenitor cells (RPCs) throughout retinogenesis, ultimately becoming restricted to Müller glia (MG) and a subset of amacrine interneurons (4, 5).

Germline deletion of *Lhx2* results in complete anophthalmia (1). However, conditional neuroretinal knockout of *Lhx2* (*Lhx*2AcKO) during later retinogenic timepoints results in premature cell cycle exit, altered RPC competence, loss of neuroretinal-derived FGFs that results in a secondary arrest in lens fiber development, and disrupted MG development (6-9). The differentiation of neurons generated following *Lhx*2ΔcKO-induced cell cycle exit appears grossly normal, though neuronal diversity is limited by RPC competence at the stage when mitotic exit occurred (6). *Lhx2* functions similarly in progenitor cells in the cerebral cortex, where it is essential for maintaining proliferative competence and developmental multipotency (10).

*Lhx2* is essential for multiple aspects of retinal gliogenesis, with early *Lhx2* loss of function resulting in RPC dropout prior to the onset of gliogenesis. *Lhx*2ΔcKO at later timepoints yields disrupted Müller differentiation, leading to morphological abnormalities and a loss of MG-specific gene expression (7, 11). Lhx2ΔcKO in fully differentiated mature MG causes cell-autonomous initiation of hypertrophic Müller gliosis in the absence of injury (4). The effect of Lhx2ΔcKO on both RPC maintenance and gliogenesis may be mediated in part by *Lhx2-* dependent activation of genes in the Notch signaling pathway. *Lhx2* is a direct transcriptional regulator of multiple Notch pathway genes in both the retina (7) and cerebral cortex (10). Notch signaling regulates the maintenance of multipotent RPCs through the downstream activation of the *Hes* family members *Hes1* and Hes5, before ultimately promoting gliogenesis through the repression of proneural bHLH genes (12-14).

The molecular mechanisms that control the pleiotropic and context-dependent functions of *Lhx2* are unclear. However, several different transcriptional co-factors function as either co-activators or co-repressors with LHX2 proteins. LIM-HD transcriptional activator function is dependent on the formation of protein complexes with LIM domain-binding (LDB) cofactors (15). Targeted loss of function of *Ldb* genes phenocopies targeted disruption of LIM-HD genes (16). Knockout of both *Ldb1* and *Ldb2* in RPCs phenocopies Lhx2ΔcKO (8), as does the misexpression of a dominant negative (DN) form of *Ldb1* in hippocampal progenitors (17). Expression of *Rnf12*, which encodes a RING finger LIM domain-interacting nuclear ubiquitin ligase, has been shown to result in the degradation of LDB proteins complexed with LIM-HD TFs, and thereby negatively regulates the transcriptional activity of LIM-HD TFs (18, 19). However, *Rnf12* has not been studied in the context of neuronal development.

In this study, we investigate the role played by *Lhx2*-interacting transcriptional coregulators during mammalian postnatal retinal development. We find that misexpression of *Lhx2*, in combination with *Ldb1*, in the neonatal mouse retina results in increased formation of rod photoreceptors at the expense of MG and bipolar interneurons, and drives a dramatic shift in amacrine cell (AC) morphology from narrow field diffuse patterns to wide field stratified patterns. We show that *Lhx2* directly regulates expression of multiple bHLH factors, and that the effects observed following misexpression are dependent on *Ascl1* and *Neurog2*, respectively. In contrast, we show that co-expression of *Rnf12* with *Lhx2* is both necessary and sufficient for Müller gliogenesis. These results identify a unique molecular switching mechanism that regulates the balance of retinal neurogenesis and gliogenesis through direct interaction with *Lhx2*, an essential master regulator of retinal development.

## Results

### Overexpression of *Lhx2* blocks Müller gliogenesis, and drives formation of rod photoreceptors and wide field amacrine cells (wfACs)

To examine the effect of misexpression of *Lhx2* on retinal development, we electroporated postnatal day (P)0 mice with control (pCAGIG) and *Lhx2-* expressing (pCAGIG-Lhx2) DNA constructs (Fig. 1a-j). *Lhx2* electroporation promoted the generation of rod photoreceptors at the expense of both MG and bipolar interneurons (Fig. 1c, d). Less than 1% of Lhx2-electroporated cells expressed either of the two MG markers P27^Kip1^ or GLUL, compared to nearly 5% of controls (Fig. 1d), and there was a significant reduction in cells with radial morphology (Fig. 1b, h-j). We also observed altered morphology among electroporated ACs (Fig. 1a, b arrows; Supplementary Fig. 1a-c). Narrow-field, diffusely arborizing ACs were generated in control electroporations (Fig. 1a), while Lhx2-electroporated RPCs generated wfACs with distinct inner plexiform layer stratification into sublamina s1, s3, and s5 (Fig. 1b, Supplementary Fig. 1a). Co-labeling of generated wfACs with AC subtype markers revealed that these cells did not express markers associated with any distinct AC subtypes, and only co-labeled with the pan-AC marker PAX6 (Supplementary Fig. 1d-m; retina in panel d also imaged at lower 40X mag in Supplementary Fig. 5b).

**Figure 1.**
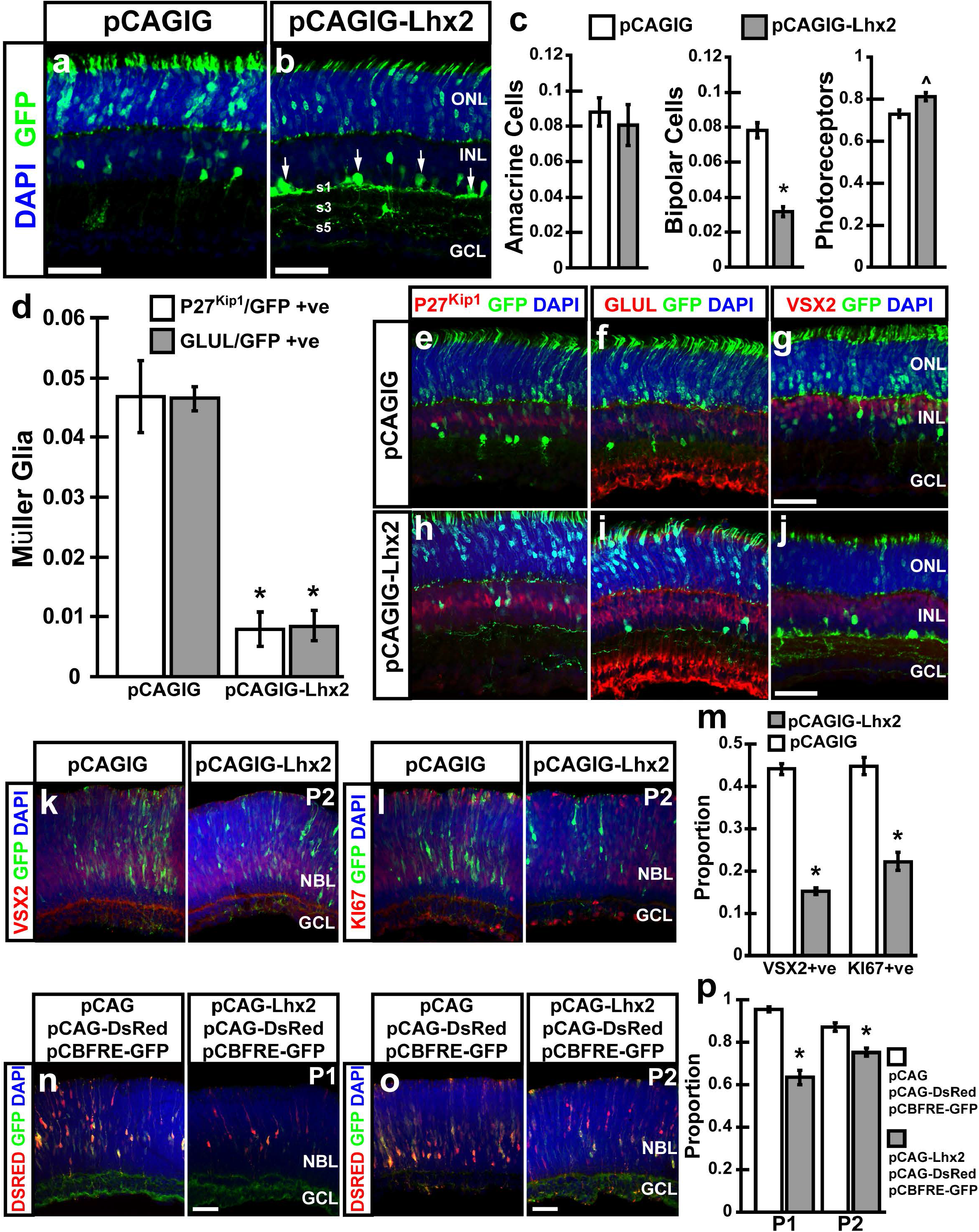
Electroporation of *Lhx2* blocks Müller gliogenesis, bipolar cell formation and changes amacrine cell morphology. (a, b, d-f, h, i) Electroporation of *Lhx2* at resulted in a significant (P<0.05) decrease at P14 of MG (P27^Kip1^ and GLUL +ve) [4.68% (SE= 0.60%, N=6, P27^Kip1^), 4.65% (SE=0.21%, N=6 GLUL) vs. 0. 8% (SE=0.29%, N=6, P27^Kip1^); 0.85% (SE=0.25%, N=6, GLUL)]. (a-c, g, j) *Lhx2* electroporation resulted in decreased (P<0.05) bipolar interneurons (VSX2 +ve) [7.81% (SE=0.38%, N=6) vs. 3.17% (SE=0.26%, N=6)] and increased photoreceptors [77.3% (SE=2.4%, N=5) vs. 82.44% (SE=2.1%, N=5)]. (b) Amacrine cell morphology changed from narrow field cells with diffuse dendrites to wide-field amacrine cells, which stratified into the S1, S2, and S3 sublamina of the inner plexiform layer (a, white arrows). (k-m) Cells electroporated with pCAGIG-Lhx2 at P0 showed significant (P<0.05) down-regulation of both VSX2 and KI67 by P2 [pCAGIG, 45.75% (SE= 2.6%, N=5, VSX2); 44.8% (SE=1.79%, N=5 KI67); pCAGIG-Lhx2, 15.3% (SE=0.42%, N=5, VSX2); 22.8% (SE=1.97%, N=5, KI67)]. (n-p) Electroporation of *Lhx2* at P0 results in a significant decrease (P<0.05) of pCBFRE-GFP Notch reporter expression at P1 and P2 [pCAG, 95.49% (SE=0.4%, N=5, P1); 87.34% (SE=1.57%, N=5, P2); pCAG-Lhx2, 63.43% (SE=2.86%, N=5, P1); 75.39% (SE=1.5%, N=5, P2)]. * Indicates statistically significant decrease. ^ Indicates statistically significant increase. GCL, ganglion cell layer; INL, inner nuclear layer; NBL, neuroblastic layer; ONL, outer nuclear layer; P, postnatal day; s inner plexiform layer sublamina. Scale bars, 50 ¼m (all panels).

### Overexpression of *Lhx2* promotes cell cycle exit and downregulation of Notch signaling

Because *Lhx2* electroporation resulted in a loss of MG and bipolar interneurons, both populations being among the last cell types generated in the retina, we tested whether *Lhx2* overexpression affected the timing of RPC cell cycle exit (Fig. 1 k-m). Electroporation of *Lhx2* resulted in premature cell cycle dropout and progenitor depletion by P2 (Fig. 1 m). The number of cells colabeled with the RPC marker VSX2 was reduced from 44% in controls to 15% in cells overexpressing *Lhx2* (Fig. 1 m). Similarly, the number of electroporated cells co-labeled with the proliferation marker KI67 was reduced from 45% in controls to 22% with *Lhx2* (Fig. 1 m).

Since *Lhx2* electroporation promoted rod photoreceptor production at the expense of bipolar cells and MG, a process that requires the inhibition of Notch signaling in newly post-mitotic retinal precursors (14), we tested whether Notch signaling was suppressed in *Lhx2* electroporated cells. P0 retinas were coelectroporated with a pCAG-DsRed cell reporter, pCBFRE-GFP Notch signaling reporter and either pCAG control or pCAG-Lhx2 construct (Fig. 1n-p). Analysis at P1 and P2 revealed significant decreases in Notch reporter labeling in cells electroporated with *Lhx2* compared to controls (95% vs. 63% at P1, p<0.05, N=5; 87% vs. 75% at P2, p<0.05, N=5) (Fig. 1p). Taken together, these results show that electroporation of *Lhx2* results in rapid cell cycle dropout and downregulation of Notch signaling.

### *Lhx2* regulates neurogenesis and neuronal differentiation in part by direct regulation of proneural and neurogenic bHLH gene expression

The phenotype resulting from misexpression of *Lhx2* closely mirrors that of *Lhx2* loss of function (7). To determine why *Lhx2* misexpression might phenocopy *Lhx2* loss of function, we first analyzed the expression of multiple proneural and neurogenic bHLH genes in an *Lhx2* ΔcKO model using the *Pdgfra-Cre; Lhx2^lox/lox^* mouse line. In these animals *Lhx2* is deleted from late-stage RPCs, resulting in a loss of MG and consequent photoreceptor degeneration (7) (Supplementary Fig. 2). RNA-Seq data obtained from *Pdgfra-Cre; Lhx2^lox/lox^* mice (7), showed substantially reduced expression, relative to controls, of multiple proneural and neurogenic bHLH genes, including *Neurod1, Neurod4, Neurog2, Ascl1, Hes6* and *Olig2* (Table ST 1). We performed *in situ* hybridization to validate these results, and found that expression of each of these genes was reduced in *Pdgfra-Cre; Lhx2^lox/lox^* mice (Fig. 2a-f; arrows). These data suggest that *Lhx2* is essential not only for expression of gliogenic bHLH factors in RPCs, as described previously (7), but also for multiple neurogenic bHLH factors. We tested this hypothesis by conducting ChlP-qPCR, examining evolutionarily conserved candidate cis-regulatory sequences located upstream of genes that contained consensus LHX2 binding sites. We found that LHX2 selectively bound to *cis-* regulatory sequences associated with the neurogenic bHLH genes *Neurod1* and *Neurod4*, as well as the proneural bHLH *Neurog2* (Fig. 2g, h). Analysis of the normalized ratio of LHX2 binding at P2 vs. P8 revealed increased occupancy at P2, correlating closely with the period of active neurogenesis (Fig. 2h). Intriguingly, the wfAC phenotype generated following *Lhx2* electroporation closely resembles phenotypes resulting from overexpression of the *NeuroD* family member *Neurod2* (20).

**Figure 2.**
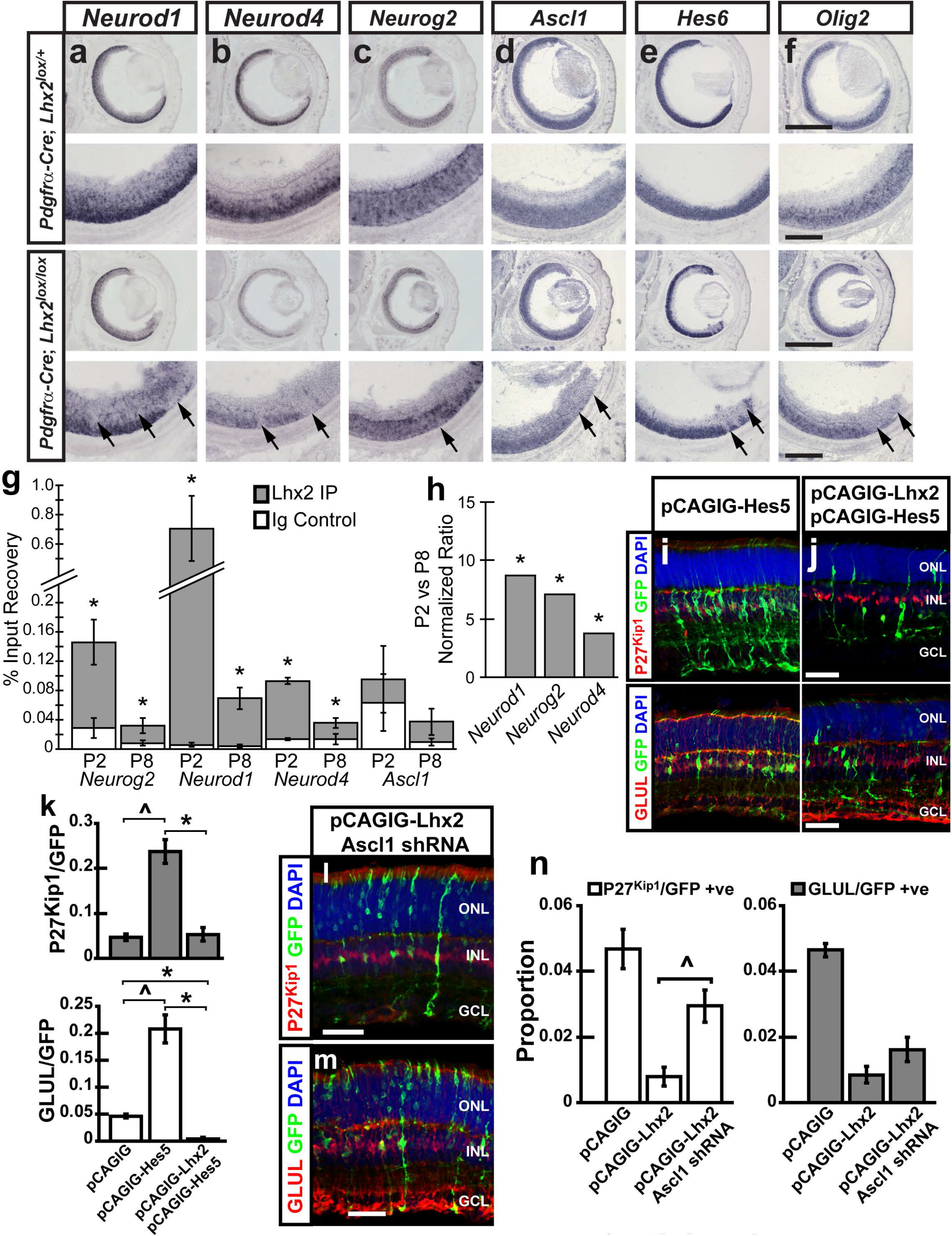
*Lhx2* regulates expression of proneural and neurogenic factors in the retina. (a-f) In situ hybridization analysis of Lhx2ΔcKO *(Pdgfra-Cre; Lhx2^o×Ao×^)* retinas at P0 reveals the requirement of *Lhx2* for proneural and neurogenic bHLH expression. Lower power images are 5X magnification, while high power images are 20X. (g) ChIP performed on retinal tissue collected at postnatal days 2 and 8. Graphs show the mean percentages of input recovery for the IP fractions and the isotype controls. * Indicates statistical significance (P<0.05). Indicated bars represent the standard error. (h) The normalized ratio of LHX2 binding to target loci reveals decreasing occupancy from P2 to P8. (i-k) Co-electroporation of *Lhx2* with *Hes5* blocked the gliogenic effect of *Hes5* electroporation [pCAGIG-Hes5, 23.75% (SE=2.63%, N=6, P27^Kip1^); 20.83% (SE=2.46%, N=6, GLUL); pCAGIG-Hes5/pCAGIG-Lhx2, 5.33% (SE= 1.4%, N=6, P27^Kip1^); 0.43% (SE=0.1%, N=6, GLUL)]. (l-n) shRNA knockdown of *AscI1* rescues (P<0.05) P27^Kip1^ expression [pCAGIG-Lhx2, 0.8% (SE=0.29%, N=6, P27^Kip1^); 0.85% (SE=0.25%, N=6, GLUL); pCAGIG-Lhx2/Ascl 1 shRNA, 2.95 % (SE=0.48%, N=6, P27^Kip1^); 1.62% (SE=0.36%, N=6, GLUL)]. * Indicates statistical significance for panels (g, h), but represents a significant decrease (k, n). ^ indicates significant increase (k, n). Scale Bars, 1000 ¼m (5 X mag, a-f), 250 ¼m (20 X mag, a-f), 50 ¼m (j, l, m).

*Hes5* encodes an E-box-selective bHLH protein which inhibits retinal neurogenesis and promotes MG specification (20). *Lhx2* function is required for the gliogenic effects of *Hes5* in retina (7). Since *Lhx2* appears to be necessary for expression of both proneural and neurogenic bHLHs, yet is essential for Hes5-dependent gliogenesis, we next tested whether simultaneous misexpression of *Lhx2* and *Hes5* could promote MG specification.

Electroporation of pCAGIG-Hes5 potently promoted the formation of MG (Fig. 2i, k). However, co-electroporation with pCAGIG-Lhx2 blocked the gliogenic effects of *Hes5*, and disrupted MG morphogenesis (Fig. 2j, k). The fraction of cells that expressed P27^Kip1^ was similar to that of vector controls, while the fraction expressing GLUL was identical to that observed in retinas electroporated with *Lhx2* alone (Fig. 2k; Fig. 1d). These data indicated that *Lhx2* expression is sufficient to override the gliogenic activity of *Hes5*, and that *Hes5* cannot suppress the neurogenic properties of *Lhx2*.

### *Lhx2* overexpression blocks gliogenesis through an ***Ascl1-***dependent mechanism, while promoting wfAC formation through ***Neurog2.***

The previously described data shows that *Lhx2* is necessary for both proneural and neurogenic bHLH expression. Furthermore, electroporation of *Lhx2* disrupts MG development, blocks Hes5-mediated MG formation, and suppresses Notch signaling in RPCs. This indicates that, although *Lhx2* is required for both Notch pathway gene expression and Notch-mediated Müller gliogenesis (7), misexpression of *Lhx2* in RPCs can inhibit Notch signaling and promote retinal neurogenesis, similar to the effects of overexpressing *Lhx2* in the hippocampus (17). One mechanism by which this might occur is through Lhx2-dependent regulation of expression of the proneural bHLH genes *Ascl1* and *Neurog2* in RPCs, which inhibit Notch (21, 22).

We first tested whether shRNA knockdown of *Ascl1* concurrent with *Lhx2* electroporation could reverse the inhibition of gliogenesis. We observed that coelectroporation of *Ascl1* shRNA constructs with pCAGIG-Lhx2 partially rescued MG differentiation, as indicated by the restoration of P27^Kip1^ positive cells that display radial morphology characteristic of MG (Fig. 2l-n). Interestingly, GLUL expression remained suppressed, indicating that *Ascl1* knockdown cannot fully rescue terminal glial differentiation.

The role of *Neurog2* is less well understood in the retina, due to its functional redundancy with *Ascl1* (22). Electroporation of pCAGIG-Neurog2 at P0 was neurogenic, resulting in the specification of PAX6+ narrow field ACs with diffuse dendritic morphology (Supplementary Fig. 3a). The population of ACs increased from 8.8% with pCAGIG to 19.8% with pCAGIG-Neurog2 (Supplementary Fig. 3f). A compensatory decrease in BCs and photoreceptors was observed in retinas electroporated with pCAGIG-Neurog2 (Supplementary Fig. 3i). Co-electroporation of pCAGIG-Neurog2 with pCAGIG-Lhx2 yielded an AC numbers that were not significantly different from controls (9.9%) (Fig. 3f). The ACs generated were wfACs with stratified dendritic morphology, a higher fraction of which co-labeled with calretinin (CALB2) (Supplementary Fig. 3b-e, i). The range of field coverage varied but was typically very large, with arbors often extending to the retinal periphery (Supplementary Fig. 3c-e). We tested whether the neurogenic and wfAC phenotypes promoted by *Lhx2* required *Neurog2* by co-electroporating pCAGIG-Lhx2 with a *Neurog2* shRNA construct. We found that knockdown of *Neurog2* expression completely blocked the formation of wfACs (Supplementary Fig. 3g, h). However, in contrast to knockdown of *Ascl1, Neurog2* knockdown did not rescue the disruption of MG development that resulted from *Lhx2* overexpression (Supplementary Fig. 3h).

**Figure 3.**
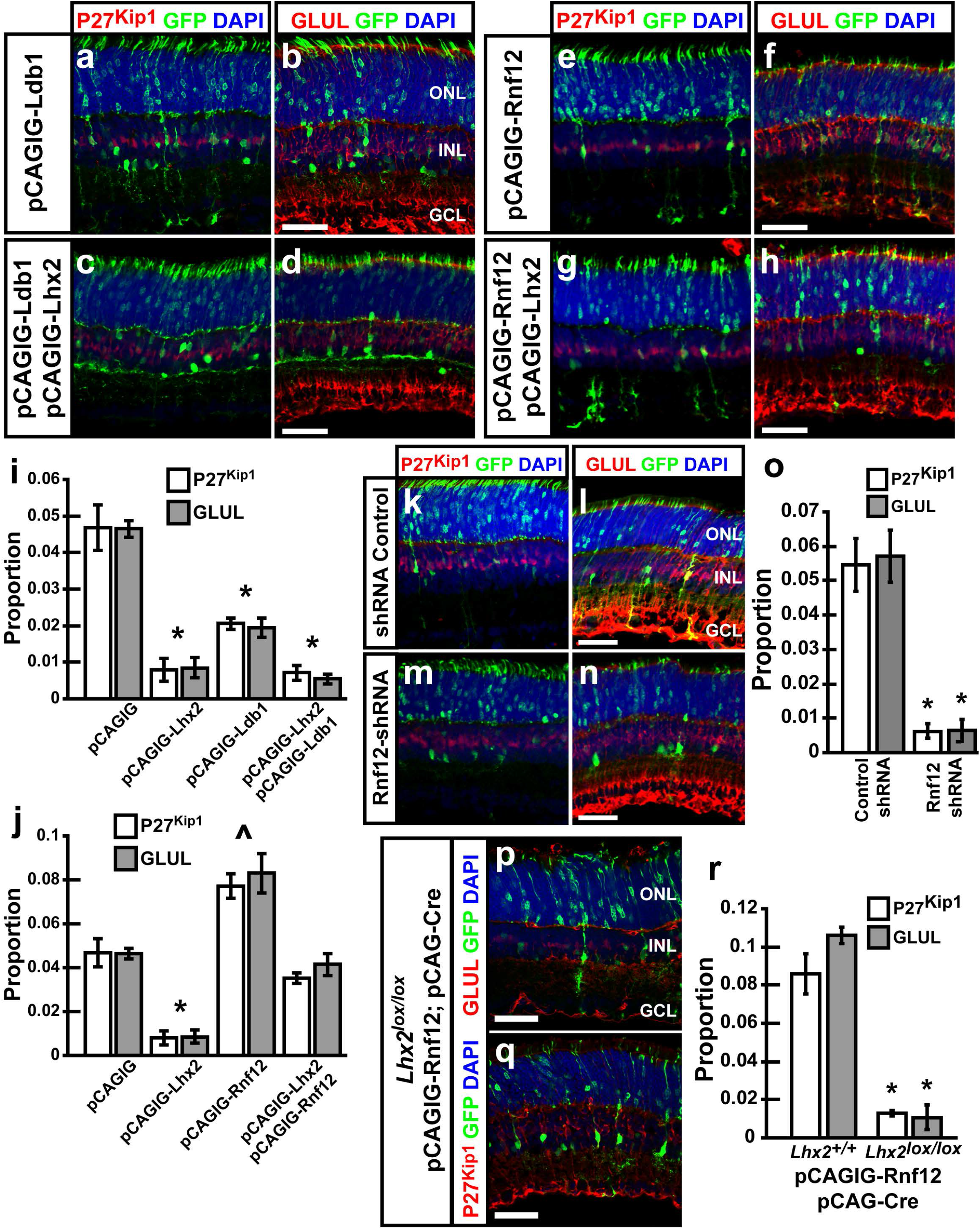
Co-electroporation of *Lhx2* with *Ldb1* or *Rnf12* differentially affects Müller gliogenesis. (a-d, i) Electroporation of *Ldb1* inhibits the formation of MG, and co-electroporation of *Lhx2* with *Ldb1* generates an identical phenotype as electroporation of *Lhx2* alone. (e-h, j) Electroporation of *Rnf12* significantly increases the proportion of MG generated [7.73% (SE=0.53%, N=6, P27^Kip1^) and 8.3% (SE=0.85%, N=6 GLUL)], while co-electroporation of *Lhx2* with *Rnf12* rescues MG [3.53% (SE=0.19%, N=6, P27^Kip1^) and 4.15% (SE=0.45%, N=6 GLUL)]. (k-o) shRNA knockdown of *Rnf12* significantly blocks the formation of MG compared to shRNA controls [5.45% (SE=0.75%, N=6, P27^Kip1^); 5.72% (SE=0.74%, N=6 GLUL) vs. 0.63% (SE=0.18%, N=6, P27^Kip1^); 0.64% (SE=0.31%, N=6, GLUL)]. (p-r) *Rnf12* requires functional *Lhx2* to promote MG development (P<0.05; N=3) P27^Kip1^, (P<0.05; N=3) GLUL. * indicates significant decrease. ^ indicates significant increase. Scale bars, 50 ¼m (all panels).

### The neurogenic role of *Lhx2* is mediated by interaction with *Ldb1*, but the *Lhx2* cofactor ***Rnf12*** activates Lhx2-dependent gliogenesis

The LIM domain binding protein LDB1 directly interacts with LHX2 (23). Recent studies of LDB function in early-stage RPCs have shown that loss of function of either *Ldb1* or *Ldb2* does not affect RPC proliferation or gliogenesis, but loss of function of both *Ldb1* and *Ldb2* genes phenocopies the loss of function of *Lhx2* (8). We found that *Ldb1* mRNA expression is broadly expressed in the developing retina, being readily detectable in RPCs in the retinal neuroblastic layer (NBL), and in differentiated neurons (Supplementary Fig. 4). Expression becomes localized primarily to ganglion cell layer (GCL) and inner nuclear layer (INL) cells in the mature retina (Supplementary Fig. 4f’’, g’’). Co-expression of *Ldb1* and *Lhx2* mRNA is observed in RPCs in the NBL from E14-P2, and in INL cells from P5 to P21 (Supplementary Fig. 4). Because of this overlap of expression of *Ldb1* and *Lhx2*, we tested whether electroporation of *Ldb1* could modify the developmental effects induced by *Lhx2* overexpression.

**Figure 4.**
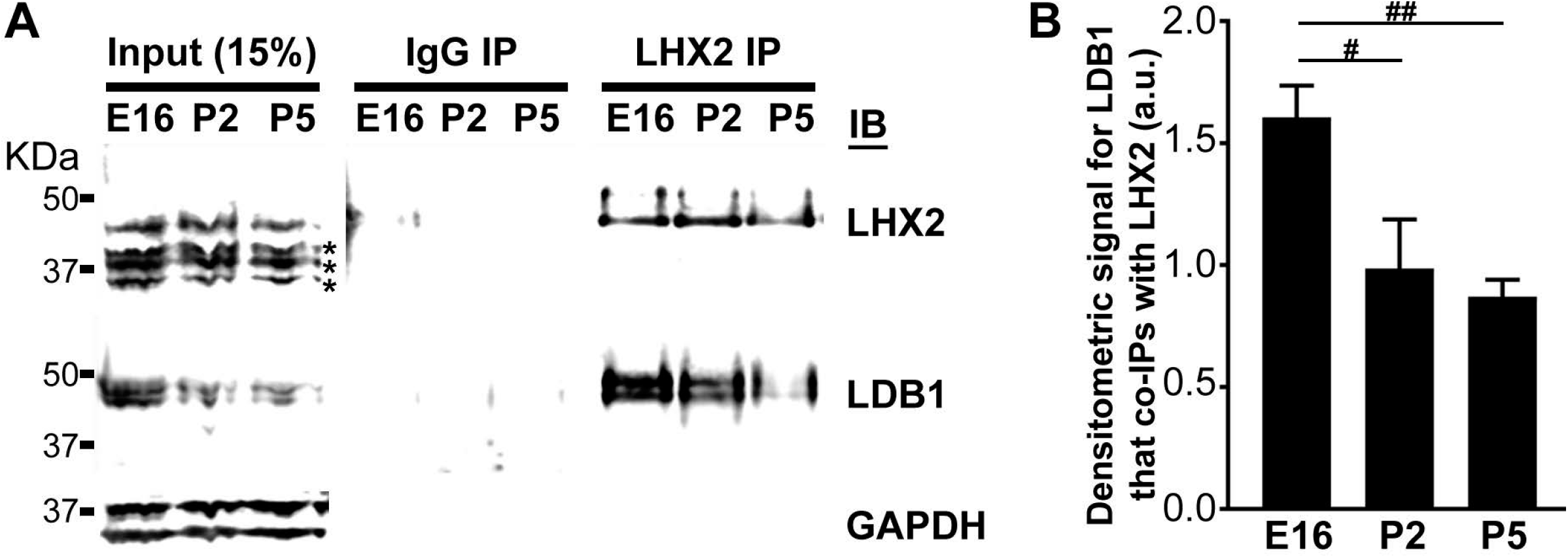
Reduced levels of the LHX2-LDB1 are seen following the onset of retinal gliogenesis. (a) Co-immunoprecipitation (co-IP) of LDB1 with LHX2 in retinal tissue collected at E16, P2 and P5. Decreased LDB1 interaction with LHX2 is observed at P5. IgG IP is used as negative control. * indicates non-specific bands detected by the anti-LHX2 antibody that do not appear in IP lanes. (b) One-way ANOVA analysis that compared the densitometry signals of LDB1 that co-IPs with LHX2 at each timepoint was found to be statistically significant (p=0.0016). All signals are normalized to Lhx2 in the age-matched input lane. Post-hoc t-test indicates that the decrease in levels of LDB1 that co-IP with LHX2 decreases significantly with E16>P2>P5. # and ## indicate statistical significance at p= 0.016 and p= 0.003 in the post-hoc t-test respectively.

Electroporation of pCAGIG-Ldb1 resulted in a significant decrease in the production of MG, from 4.7% P27^Kip1^+ve and 4.7% GLUL+ve in control electroporated cells, to 2.1% and 2.0%, respectively (Fig. 3a, b, i). The reduction was less pronounced than that observed following electroporation with pCAGIG-Lhx2, and unlike *Lhx2* electroporation no notable changes in AC morphology were observed (Fig. 3a, b; Supplementary Fig. 5). Outside of the reduction in MG, no significant changes in the patterns or morphology of electroporated cells could be distinguished between pCAGIG and pCAGIG-Ldb1 (Fig. 3a, b; Supplementary Fig. 5). Co-electroporation of pCAGIG-Lhx2 with pCAGIG-Ldb1 produced a phenotype identical to that observed in *Lhx2* electroporations - a significant loss of MG and production of wfACs (Fig. 3c, d, i; Supplementary Fig. 5).

We also investigated the effects of loss of LDB function by overexpressing a dominant-negative (DN) construct of Ldb1, which has previously been shown to phenocopy loss of *Lhx2* function in hippocampal progenitors (17). We observed that overexpression of DN-LDB1 in P0 retina phenocopies the previously described loss of function of *Lhx2* (7), resulting in a loss of MG, but not in the fraction of BCs or photoreceptors (Supplementary Fig. 6). These results confirmed that LDB factors are indeed necessary for Lhx2-mediated regulation of RPC maintenance and gliogenesis.

Since developmental outcomes mediated by LIM-HD factors, including LHX2, are co-regulated by *Rnf12* (18, 19), we tested whether *Rnf12* expression could alter *Lhx2* function in the retina. Analysis of *Rnf12* mRNA expression in the developing retina revealed relatively low expression in RPCs at from E14-E18 timepoints (Supplementary Fig. 4a’-c’). Postnatal expression revealed a distinct upregulation and enrichment of RNA expression in subsets of cells in the NBL from P0 to P2 and in the medial INL at P5, consistent with the spatial and temporal onset of Müller gliogenesis (Supplementary Fig. 4d’-f’), as well as with previous studies which reported increased expression of *Rnf12* in MG precursors (24).

Electroporation of pCAGIG-Rnf12 at P0 led to a significant increase in the production of MG from 4.7% and 4.6% (P27^Kip1^+ve and GLUL+ve respectively) in controls, to 7.72% and 8.3 % (Fig. 3e, f, j). Furthermore, co-electroporation of *Rnf12* with *Lhx2* rescued the reduction in gliogenesis observed following electroporation of *Lhx2* alone (Fig. 3g, h, j). We also observed that coelectroporation of *Rnf12* with *Lhx2* reversed the observed changes in amacrine cell morphology that resulted from *Lhx2* electroporation, preventing the formation of wfACs (Fig. 3g, h; Supplementary Fig. 5f). Electroporation of *Rnf12* alone or with *Lhx2* inhibited the formation of ACs broadly (Supplemental Fig. 5e-g). We next tested whether *Rnf12* was required for glial development. Electroporation with shRNA constructs targeting *Rnf12* at P0 resulted in a loss of MG as determined by P27^Kip1^ and GLUL immunostaining (Fig. 3k-o). The relative loss of MG was nearly identical to that reported following *Lhx2* loss of function (7). To determine if *Rnf12* requires *Lhx2* in order to promote MG differentiation, we coelectroporated pCAGIG-Rnf12 with pCAG-Cre into *Lhx2+^/^+* and *Lhx2^lox/lox^* retinas at P0. Concurrent loss of function of *Lhx2* blocked the Rnf12-dependent increase in gliogenesis (Fig. 3p-r). In these mice, the proportion of P27^Kip1^ and GLUL+ve electroporated cells (1.3% and 1.1% respectively) was nearly identical to that reported following *Lhx2* loss of function (7). Taken together, these results suggest that *Rnf12* acts through an Lhx2-dependent mechanism in late-stage RPCs to induce gliogenesis.

### Lower relative levels of LHX2-LDB1 protein complexes are seen as gliogenesis is initiated

We have previously shown that LHX2 interacts with LDB1 in developing retina at E15.5 and P0.5 (8). The findings in this study, suggest that upregulation of *Rnf12* may lead to the relative fraction of LHX2 bound to LDB1 decreasing during gliogenesis due to RNF12-dependent degradation of LDB1. To test this hypothesis, we performed immunoprecipitation analysis of LHX2 and LDB1 at three different timepoints. We analyzed LHX2 protein complexes at E16, when only neurons are born; P2, when gliogenesis is beginning; and P5, when gliogenesis peaks (25). Both LHX2 and LDB1 were expressed and directly interacted with one another at all three stages (Fig. 4a). However, when immunoprecipitation was performed with antibodies to LHX2, levels of recovered LDB1 levels showed a substantial reduction at P2 relative to E16, and even more pronounced reductions at P5 (Figure 4b), when normalized to total levels of immunoprecipitated LHX2.

## Discussion

The molecular mechanisms that control CNS gliogenesis are still poorly understood. Work from many groups has shown that a combination of genes encoding extrinsic and intrinsic signals control gliogenesis. Extrinsic signals include the Notch/Delta pathway, while several transcription factors - including *Sox9, Nfia, Hes5* and *Zbtb20* - have been shown to be either necessary or sufficient to induce gliogenesis (20, 26, 27).

Here, we shed light on the mechanism by which context-specific functions of *Lhx2* are regulated during retinal development. LHX2 regulates expression of multiple different Notch pathway genes (7), and this is of central importance to its role in actively regulating the balance of neurogenesis and gliogenesis in RPCs. Furthermore, LHX2 directly activates expression of Notch-regulated bHLH factors - such as *Ascl1, Neurog2* and *Hes5*, as well as multiple Notch-independent neurogenic bHLH factors such as members of the *NeuroD* family (28, 29), which in turn induce wfAC formation when overexpressed (30). In this study, we confirm and extend previous work that demonstrated an essential role for *Ldb1-Lhx2* function in both RPC proliferation and retinal gliogenesis. We show that ectopic *Lhx2* expression potently suppresses Notch signaling, resulting in early RPC cell cycle exit and blocking retinal gliogenesis, while at the same time promoting formation of wfACs. We further demonstrate that these effects are dependent on Lhx2-dependent regulation of the expression of multiple proneural bHLH factors. We also show that *Rnf12*, which specifically ubiquitinates LDB proteins and targets them for proteolysis, is both necessary and sufficient to promote Müller glial development, and does so in a strictly Lhx2-dependent manner. Upregulation of *Rnf12* expression correlated with a reduction in retinal levels of the LHX2-LDB1 complex, concurrent with the onset and progression of gliogenesis. A model summarizing these findings, and integrating them with previous studies of *Lhx2* function in postnatal retinal development, is shown in Figure S7.

These findings highlight the importance of LIM cofactor-mediated Lhx2-dependent transcriptional activation in controlling cell fate specification in the CNS. This does not, however, exclude a parallel function of *Rnf12* in promoting Lhx2-dependent transcriptional repression. While Lhx2-dependent transcriptional activation is dependent on LDB proteins, Lhx2-dependent transcriptional repression involves recruitment of histone modifying enzymes, including the HDAC and NuRD protein complexes (23, 31). RNF12 itself can directly bind both LHX2 and SIN3A, leading to recruitment of HDAC proteins (23). It is thus possible that *Rnf12* may promote gliogenesis by both attenuating *Lhx2-Ldb1*-dependent activation of proneural genes, and by triggering LHX2-dependent repression of these genes. Further studies will be needed to resolve this issue.

In both the retina and other CNS regions, *Lhx2* acts as a selector gene that simultaneously activates and represses different sets of tissue and/or state-specific genes. In early-stage RPCs, *Lhx2* simultaneously activates expression of RPC-specific genes, while suppressing genes that are specific to anterodorsal hypothalamus and thalamic eminence (32). Likewise, in early cortical progenitors, *Lhx2* activates cortical plate-specific genes, while repressing expression of genes enriched within the cortical hem (33). The dynamic regulation of *Rnf12* and *Ldb* activity, which plays a central role in control of retinal cell fate, may also be important for these selector functions of *Lhx2*.

## Materials and Methods

### Animals

Timed pregnant CD-1 mice were purchased from Charles River Laboratories. *Lhx2^lox/lox^* mice were bred and genotyped as previously described (7). All experimental procedures were pre-approved by the Institutional Animal Care and Use Committee of the Johns Hopkins University School of Medicine.

### Electroporation, immunohistochemistry, *in situ* hybridization and cell counts

Retinas were electroporated at P0 as previously described (7, 11) and harvested for analysis at P1, P2, or P14. *In situ* hybridization was performed as previously described (34). Immunohistochemistry and cell counting was performed as previously described (4, 7).

### ChIP-qPCR, immunoprecipitation and immunoblotting

ChIP-qPCR was performed as previously described (7). Immunoprecipitation and immunoblotting were performed as described (35).

A detailed description of all protocols and reagents used can be found in the Supplemental Information.

## Acknowledgements

We thank K. Yang for help with statistical analysis and W. Yap for comments on the manuscript. This study was supported by grants from the NIH to B.S.C. (F32EY024201, K99EY027844) and S B. (R01EY020560).

## Materials and Methods

### Animals

Timed pregnant CD-1 mice used for *in situ* hybridization, electroporation and ChIP were purchased from Charles River Laboratories. Mice were housed in a climate-controlled pathogen free facility, on a 12 hour-12 hour light/dark cycle (08:00 lights on-20:00 lights off). *Pdgfra-Cre* (stock #013148) mice were purchased from the Jackson Laboratory while *Lhx2^lox/lox^* mice were obtained from Dr. Edwin Monuki, University of California, Irvine. *Lhx2^lox/lox^; Pdgfra-Cre* and *Lhx2^+/+^; Pdgfra-Cre* mice were bred and maintained in the lab as previously described (1). All experimental procedures were preapproved by the Institutional Animal Care and Use Committee of the Johns Hopkins University School of Medicine.

### Cell counts

All counts were performed blinded on whole retinal sections or dissociated retinas as previously described (1, 2). Differences between the two means were assessed using an unpaired two-tailed Student’s t-test.

### Chromatin immunoprecipitation (ChIP)

CD-1 mice were sacrificed at postnatal day (P)2 and P8 according to Johns Hopkins IACUC animal policies. ChIP was performed as previously described (1). Whole dissected retinas were dissociated in a collagenase I suspension, cross-linked in 1% formaldehyde, quenched in 125 mM glycine and the extracted nuclei were sheared to produce 100 to 500 bp fragments by means of probe sonication. Chromatin was immunoprecipitated by using goat anti-Lhx2 antibody (Santa Cruz Biotechnology) or the related isotype control (Abcam), retained on agarose beads (Invitrogen), washed and purified by organic extraction.

Candidate target genes demonstrating altered expression levels in *Lhx2* conditional knockout retinas by RNA-Seq were screened for LHX2 consensus binding sites within annotated regulatory regions by querying the JASPAR repository database (3), and was based on GSE48068 (4). Computationally inferred Lhx2 binding sites and proximal negative control regions were analyzed in ChIP-enriched fractions and isotype controls by SYBR-qPCR (Agilent Technologies).

### Electroporation

Retinas were electroporated at P0 as previously described, and harvested for analysis at P1, P2, or P14 subject to the requirements of the study. DNA constructs used for gene misexpression in this study are as follows: pCAGIG (Addgene plasmid 11159, deposited by C. Cepko and modified into a Gateway destination vector in lab), pCAGIG-Hes5 (Gateway cloned from Ultimate Human ORF Collection (Life Technologies)), pCAGIG-Ldb1 (Gateway cloned from Ultimate Human ORF Collection (Life Technologies)), pCAGIG-Lhx2 (Gateway cloned from Ultimate Human ORF Collection (Life Technologies)), pCAGIG-Ngn2 (NeuroG2) (Gateway cloned from Ultimate Human ORF Collection (Life Technologies)), pCAGIG-Rnf12 (Gateway cloned from Ultimate Human ORF

Collection (Life Technologies)). DNA constructs used for Notch reporter analysis in this study are as follows: pCAG (modified from pCAGIG), pCAG-DsRed (Addgene plasmid 11151, deposited by C. Cepko), pCAG-Lhx2 (Gateway cloning from Ultimate Human ORF Collection (Life Technologies)), pCBFRE-GFP (Addgene plasmid 17705, deposited by N. Gaiano). DNA constructs used for shRNA knockdown in this study are as follows: Ascl1 shRNA (clone TRCN0000075398, TRC-Open Biosystems), Rnf12 shRNA (clone TRCN0000095740, TRC-Open Biosystems), Ngn2 (Neurog2) shRNA (clone FP-301 obtained from Franck Polleux, Columbia University) (5, 6), (Control (pLKO.1 vector control, TRC-Open Biosystems). All shRNA constructs have been previously shown to give substantial (>70%) knockdown of their target gene.

DNA constructs used for Lhx2 loss of function in this study are as follows: pCAG-Cre (Addgene plasmid 13775, deposited by C. Cepko), and pCALNL-GFP (Addgene plasmid 13770, deposited by C. Cepko).

### Immunohistochemistry

Antibodies utilized for fluorescent immunohistochemistry are as follows: goat anti-Brn3 (1:200; Santa Cruz Biotechnology), mouse anti-calbindin (Calb1) (1:200; Sigma-Aldrich), rabbit anti-Calretinin (Calb2) (1:200; Chemicon), goat anti-Chat (1:100; Chemicon), sheep anti-Chx10 (Vsx2) (1:200; Exalpha Biologicals), rabbit anti-Dab1 (1:200; EMD Millipore), rabbit anti-DsRed (1:500; Clontech Laboratories), rabbit anti-GABA (1:200; Sigma), mouse anti-Gad6 (Gad2) (1:200; Developmental Studies Hybridoma Bank, University of Iowa), goat anti-GFP (1:500; Rockland Immunochemicals), rabbit anti-GFP (1:1000; Invitrogen), mouse anti-Glutamine synthase (Glul) (1:200; BD Biosciences), rat anti-Glycine (1:200; ImmunoSolution), mouse anti-Islet1 (1:200; Developmental Studies Hybridoma Bank), mouse anti-Ki67 (1:200; BD Biosciences), rabbit anti-Lhx2 (1:1500; generated in house with Covance), mouse anti-P27 (1:200; Invitrogen), mouse anti-Pax6 (1:200; Developmental Studies Hybridoma Bank), rabbit anti-TH (1:500; Pel Freez), mouse anti-VGlut3 (1:200; Antibodies Incorporated). Secondary antibodies used were FITC conjugated donkey antigoat IgG (1:500; Jackson Immunoresearch), FITC conjugated donkey anti-mouse IgG (1:500; Jackson Immunoresearch), FITC conjugated donkey anti-rabbit IgG (1:500; Jackson Immunoresearch), Texas Red conjugated donkey anti-goat IgG (1:500; Jackson Immunoresearch), Texas Red conjugated donkey anti-mouse IgG (1:500; Jackson Immunoresearch), Texas Red conjugated donkey anti-rabbit IgG (1:500; Jackson Immunoresearch), Texas Red conjugated donkey anti-sheep IgG (1:500; Jackson Immunoresearch). All section immunohistochemical data shown was imaged and photographed on a Zeiss Meta 510 LSM confocal microscope.

### *In Situ* Hybridization

Single-color *in situ* hybridization was performed as previously described (7). RNA probes were generated using the following EST sequences as templates: *AscI1*, GenBank accession number BE953927; *Hes6*, GenBank accession number AW048812; *Neurod1*, GenBank accession number AI835157; *Neurod4*, GenBank accession number AI846749, *Neurog2*, GenBank accession number BC055743; *Olig2*, GenBank accession number AI844033.

### Immunoblotting and Immunoprecipitation

Wildtype retinal tissues were harvested from E16 (8 litters), P2 (5 litters) and P5 (5 litters), and snap-frozen for storage. After pooling tissues from all litters, tissue homogenization was carried out by aspirating the tissue 20 times using a 23-gauge needle in lysis buffer (100mM Tris-HCl, 150mM NaCl, 25mM NaF,50 ¼M ZnCl_2_, 15% glycerol, 1% Triton X-100) supplemented with protease inhibitors (Roche #11697498001) and BitNuclease (Biotools #B16003) for clarification. Following a 1hr of incubation at 4°C, supernatant were collected after centrifuging at 10,000 rcf for 10mins at 4°C. Following normalization using standard BCA assay, immunoprecipitation (IP) was carried out by first incubating lysates overnight at 4°C with 5ug of anti-LHX2 (clone ID: R911. 1.2E3, CDI labs Inc. catalog #15-389) and mouse pan-IgG (#sc-2025, Santa Cruz) respectively. Next, the antibody-protein complexes were pulled down by incubating 2 hours with ProteinG Dynabeads (ThermoFisher #10004D) at 4°C, washed thrice with lysis buffer and eluted in LDS-sample loading buffer (ThermoFisher #NP0008). Input lysate along with the IP samples were resolved in a SDS-PAGE gel and immunoblotted sequentially using anti-LHX2 (1:750, clone ID: R911. 1.2E3, CDI labs Inc. catalog #15-389), LDB1 (1:1000 Sigma #HPA034488), GAPDH (Sigma #G8795) and RLIM (1:2000, Millipore #ABE1949) antibodies. Anti-rabbit IRDye680RD (LiCor # 925-68071) and light-chain specific anti-mouse AlexaFluor 790 (Jackson Immunolabs #115-655-174) secondary antibodies were used to visualize bands, and blots were imaged using an infra-red fluorescence imager (LiCor Clx).

### Densitometry and Statistical Analysis

Three technical repeats of the co-IP experiments were performed, followed by three independent immunoblots. Densitometry signal from lanes corresponding to LHX2 input, LHX2 IP, LDB1 IP and GADPH loading controls of all three SDS-PAGE gels was measured using LiCor image studio software. Following normalization of signal from LHX2 IP to its respective input, the ratio of LDB1 co-IP with LHX2 was calculated. Statistical analysis was performed using R software. We performed linear regression (r, lm) to adjust batch effects in the ratio of LDB1 co-IP with LHX2 between the three blots. Next, we performed one-way ANOVA (r, aov) using the adjusted values to test if there were any statistically significant differences between the means of LDB1 signal that co-IPed with LHX2 in the E16, P2 and P5 samples. For post-hoc pairwise comparisons, we performed a t-test (r, t-test).

## Supplemental Figures

**Supplemental Figure 1.**
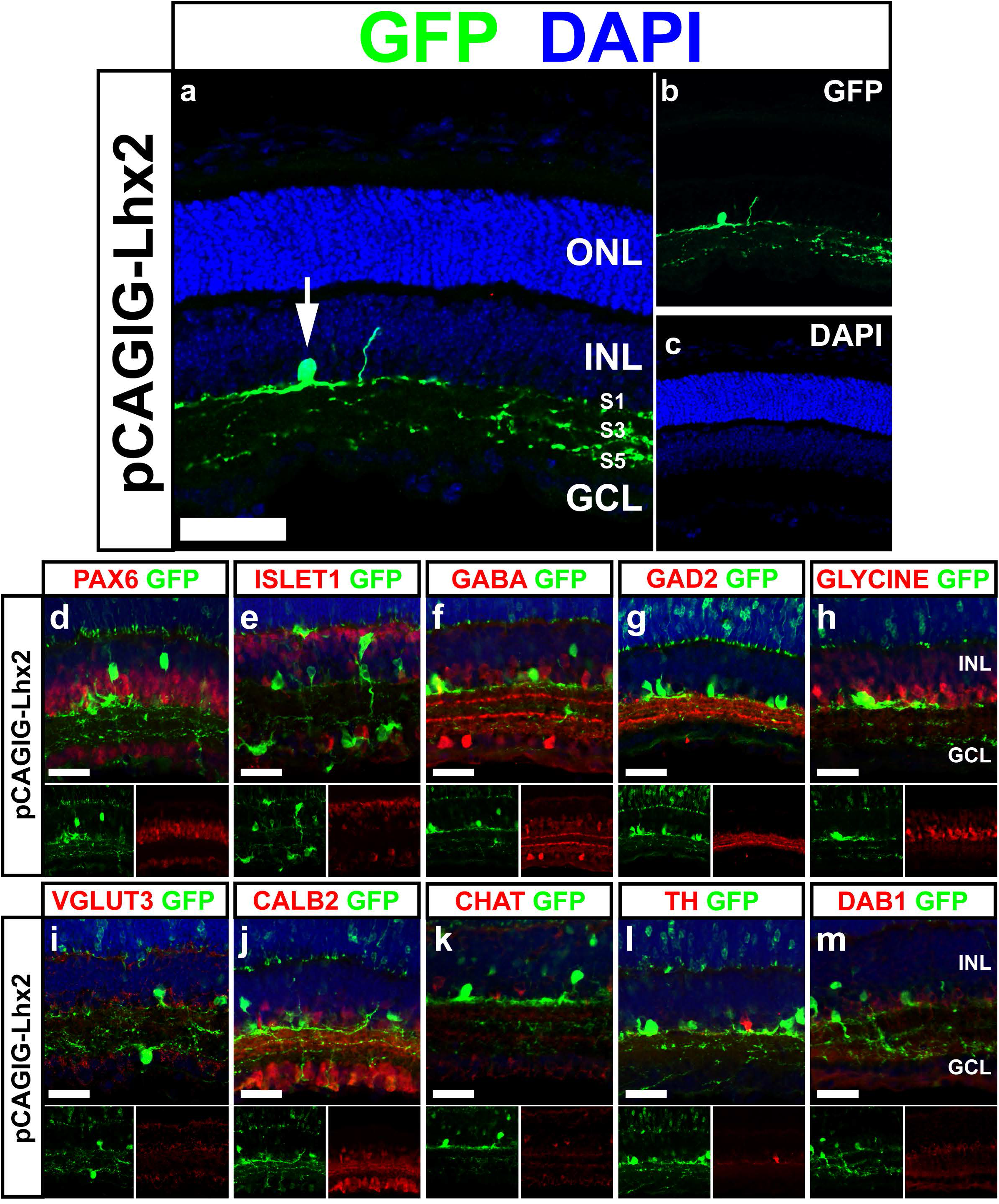
Electroporation of *Lhx2* promotes the formation of wfACs. (a-c) Morphology of a wfAC generated following electroporation of *Lhx2*. (d) Generated wide field amacrine cells co-express the pan-amacrine marker PAX6. (e-m) Co-labeling with amacrine cell subtype selective markers reveals that amacrine cells generated by *Lhx2* electroporation do not fall within any well-established molecular category. ISLET 1, CHAT-cholinergic starburst amacrine cells; GABA, GAD2-GABAergic amacrine cells; GLYCINE-glycinergic amacrine cells; VGLUT3-glutamatergic amacrine cells; CALB2-mixed population primarily AII amacrine cells, A19 amacrine cells, and non-AII glycine immunoreactive amacrine cells; TH-dopaminergic wide field amacrine cells; DAB1-AII amacrine cells. GCL, ganglion cell layer; INL, inner nuclear layer; outer nuclear layer; s inner plexiform layer sublamina. Scale bars, 50 ¼m (all panels).

**Supplemental Figure 2.**
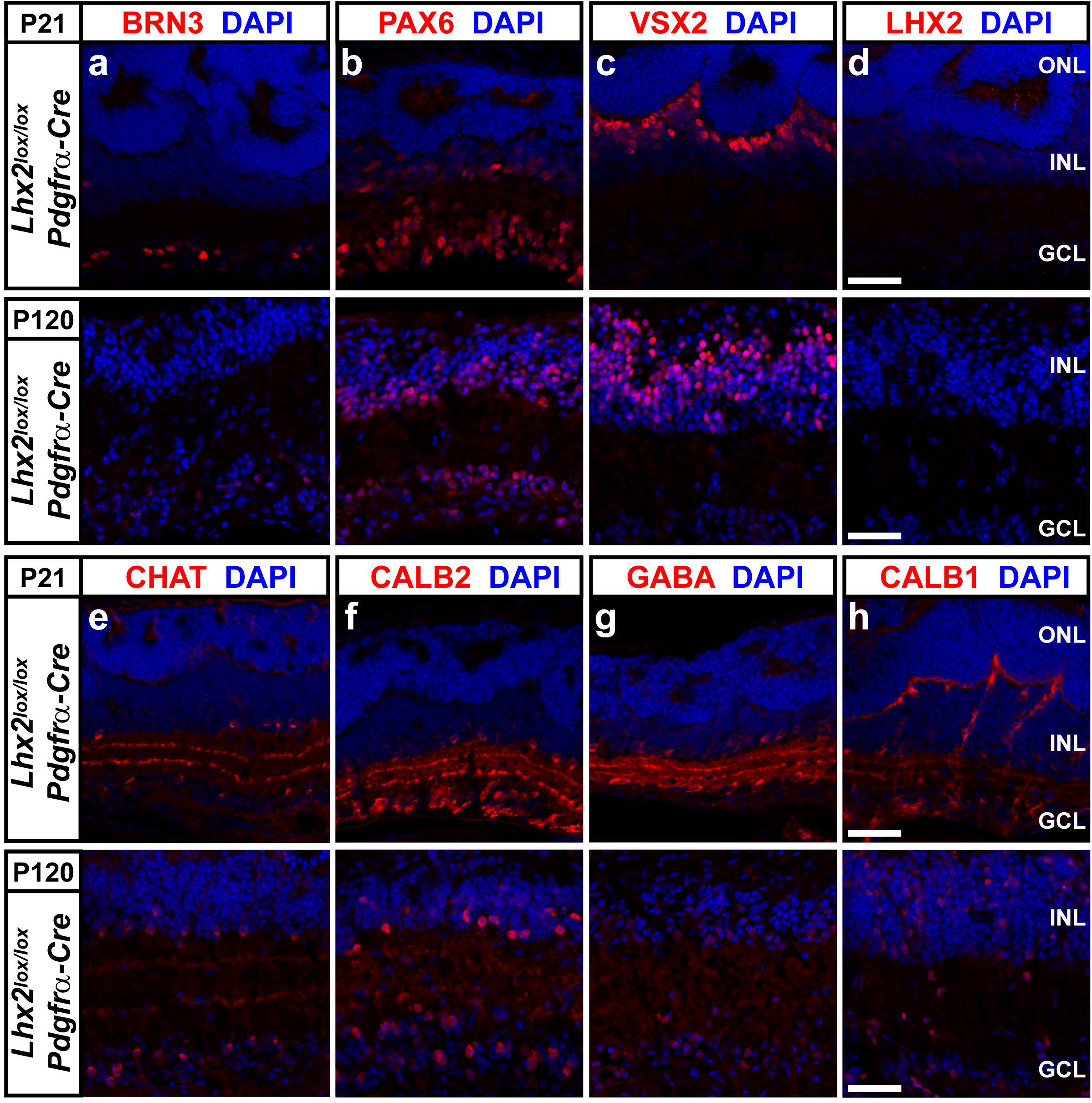
(a-d) Expression of inner retinal cell class markers at P21 and P120 in *Pdgfra-Cre; R26YFP; Lhx2^ox/lox^* retinas. Expression of the retinal ganglion cell marker Brn3 (a), retinal ganglion, amacrine, and horizontal cell marker Pax6 (b), and bipolar cell marker Vsx2 (c) are detectable at both P21 and P120. (d) Expression of Lhx2 is not detectable at both P21 and P120. (e-h) expression of amacrine cell subclass specific markers at P21 and P120 in *Pdgfra-Cre; R26YFP; Lhx2^lox/lox^* retinas. Expression of choline acetyltransferase, Chat (e), calretinin, Calb2 (f), GABA, (g), and calbindin, Calb1 (h) are detectable at both P21 and P120. Scale bars, 50 ¼m (d, h).

**Supplemental Figure 3.**
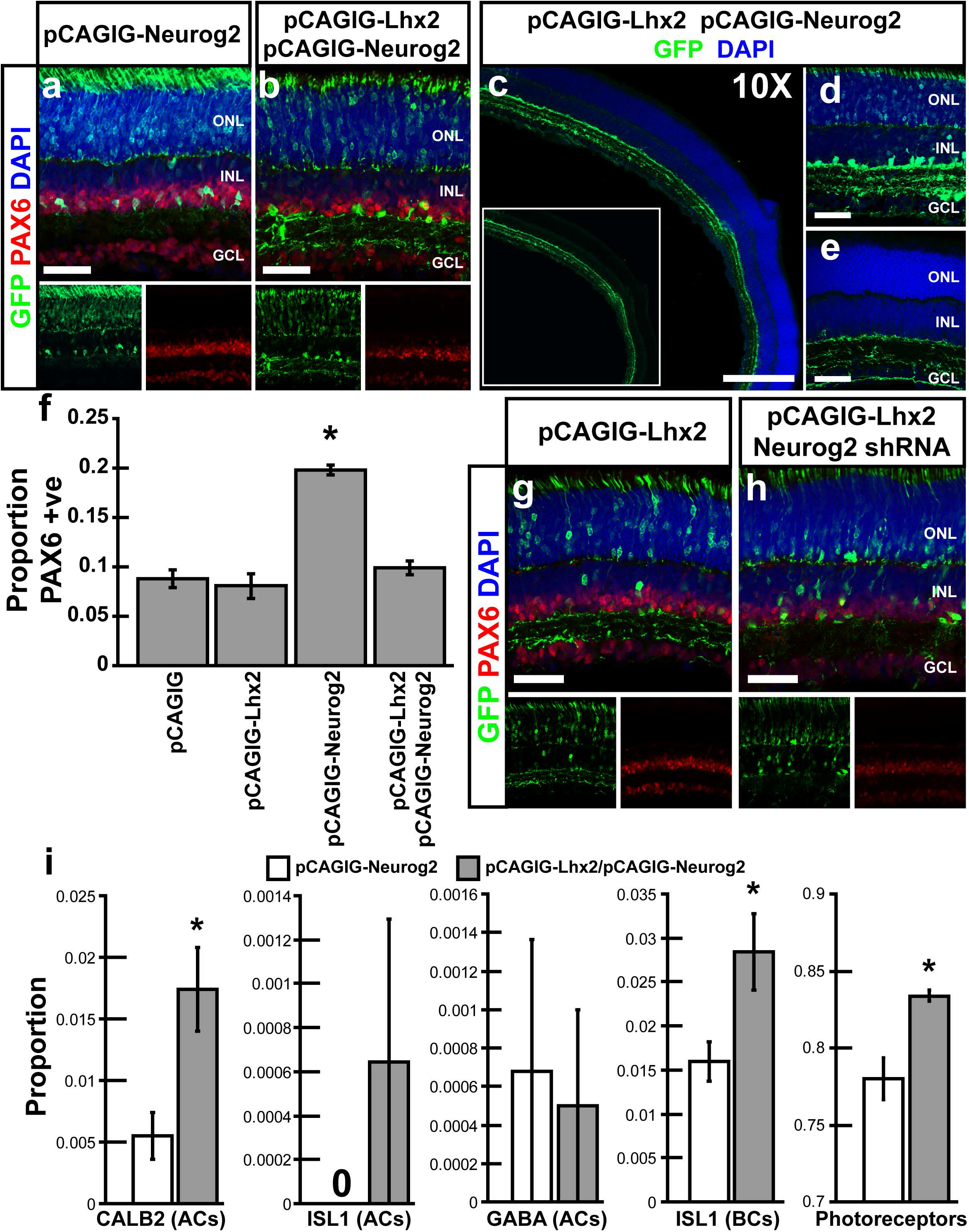
*Lhx2* synergistically promotes the formation of wide field amacrine cells with *Neurog2*. (a, f) Electroporation of *Neurog2* results in an increase in the formation of narrow field diffusely arborizing amacrine cells. (b, f) Co-electroporation of *Lhx2* with *Neurog2* transforms the morphology of amacrine cells from narrow field and diffusely arborizing to wide field and selectively stratified. The overall fraction of amacrine cells, however, is unchanged relative to that seen following electroporation of *Lhx2* alone. (c-e) Electroporation of *Lhx2* with *Neurog2* results in a synergistic expansion of the width of the dendritic field. (g, h) shRNA mediated knockdown of *Neurog2* blocks the formation of wide field amacrine cells generated by electroporation of *Lhx2*. (i) Coelectroporation of *Lhx2* with *Neurog2* results in significant increases in CALB2+ amacrine cells (primarily AII), bipolar cells, and photoreceptors compared to electroporation of *Neurog2* alone (P<0.05). Scale bars, 50 ¼m (a, b, d, e, g, h), 200 ¼m (c).

**Supplemental Figure 4.**
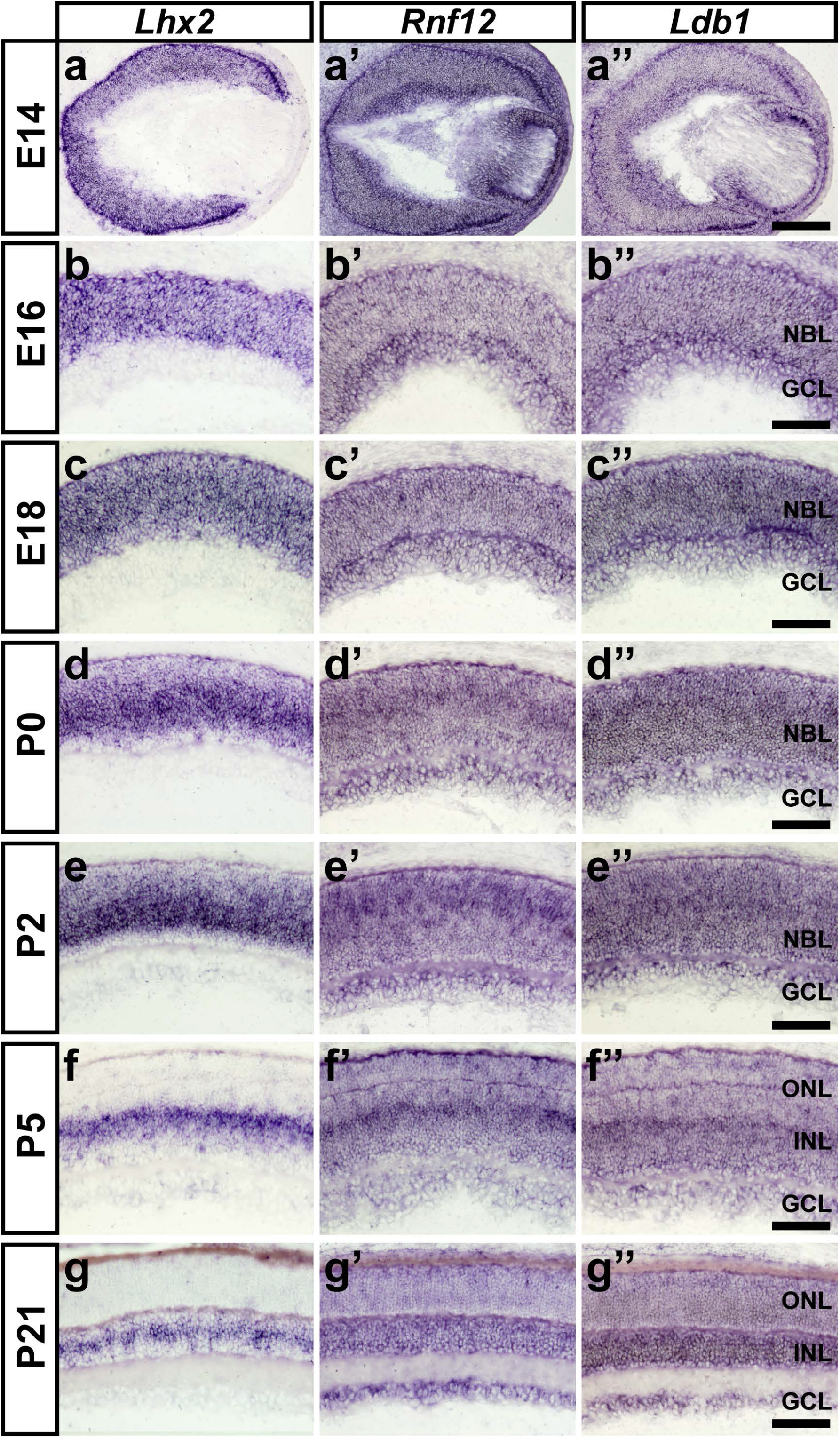
RNA expression of *Rnf12* and *Ldb1* contrasted with *Lhx2* during mouse retinal development. (a-c) RNA expression of *Lhx2* is restricted to RPCs during embryonic time points with down regulation occurring in early-born neurons in the GCL. (d-g) Down regulation of *Lhx2* in newly generated neurons continues in postnatal retina, with *Lhx2* becoming restricted to MG and subsets of amacrine cells consistent with previous reports. (a’-c’) Low levels of *Rnf12* RNA expression are seen in the NBL during embryonic time points, with higher expression detected in the GCL at E14 and E18. (d’-f’) *Rnf12* expression is up regulated in the medial NBL at P0 and remains robustly expressed in the NBL and medial INL at P2 and P5. (g’) Adult expression of *Rnf12* is located in all three retinal layers, with the INL showing the strongest labeling. (a’’-c’’) *Ldb1* expression was detected throughout the embryonic retina. (d’’-e’’) Neonatal expression of *Ldb1* is enriched in the NBL. (f’’) At P5 enrichment of *Ldb1* in the medial INL is detected. (g’’) Adult expression of *Ldb1* is localized in the INL and subsets of cells in the GCL. Weaker expression of *Ldb1* was also detected in the ONL. Scale bars, 200 ¼m (a’’), 100 um (b’’-g’’).

**Supplemental Figure 5.**
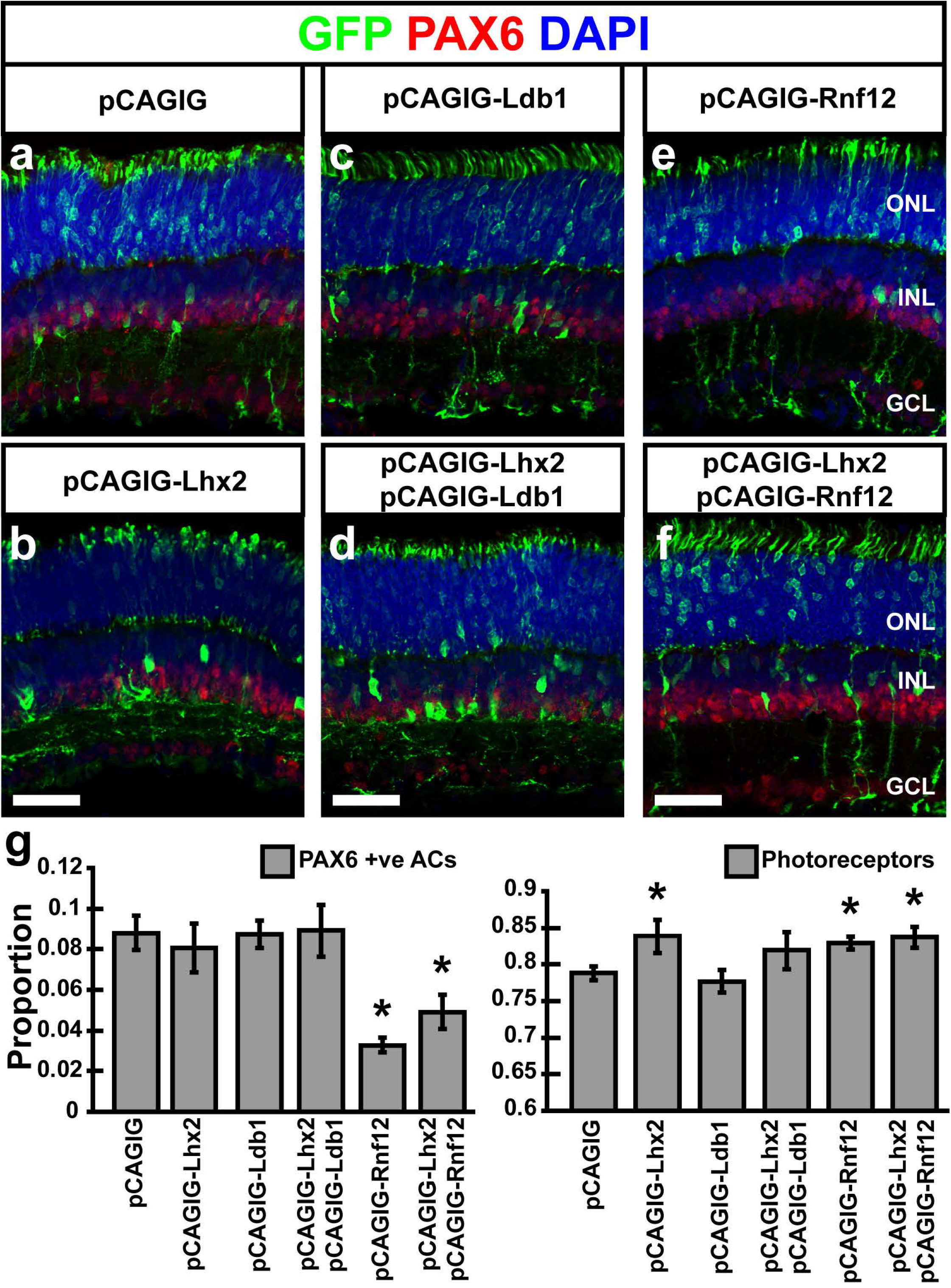
Co-electroporation of *Lhx2* with *Ldb1* or *Rnf12* results in changes in neurogenesis. (a-c, g) electroporation of *Lhx2* or *Ldb1* does not alter the proportion of amacrine cells (PAX6 +ve) generated. (d, g) Coelectroporation of *Lhx2* with *Ldb1* generates an identical wide field amacrine cell phenotype as electroporation of *Lhx2* alone. (e-g) Electroporation of *Rnf12* inhibits the formation of amacrine cells, while co-electroporation of *Rnf12* with *Lhx2* blocks the formation of wide-field amacrine cells generated by electroporation of *Lhx2* alone (P<0.05; N=6; PAX6 +ve, pCAGIG-Rnf12 vs. pCAGIG; (P<0.05; N=6; PAX6 +ve, pCAGIG-Rnf12/Lhx2 vs. pCAGIG). (b, e-g) Electroporation of Lhx2, Rnf12, or Lhx2 and Rnf12 results in mild increases in photoreceptor numbers (P<0.05; N=6). *, indicates significant decrease. Scale bars, 50 ¼m (all panels).

**Supplemental Figure 6.**
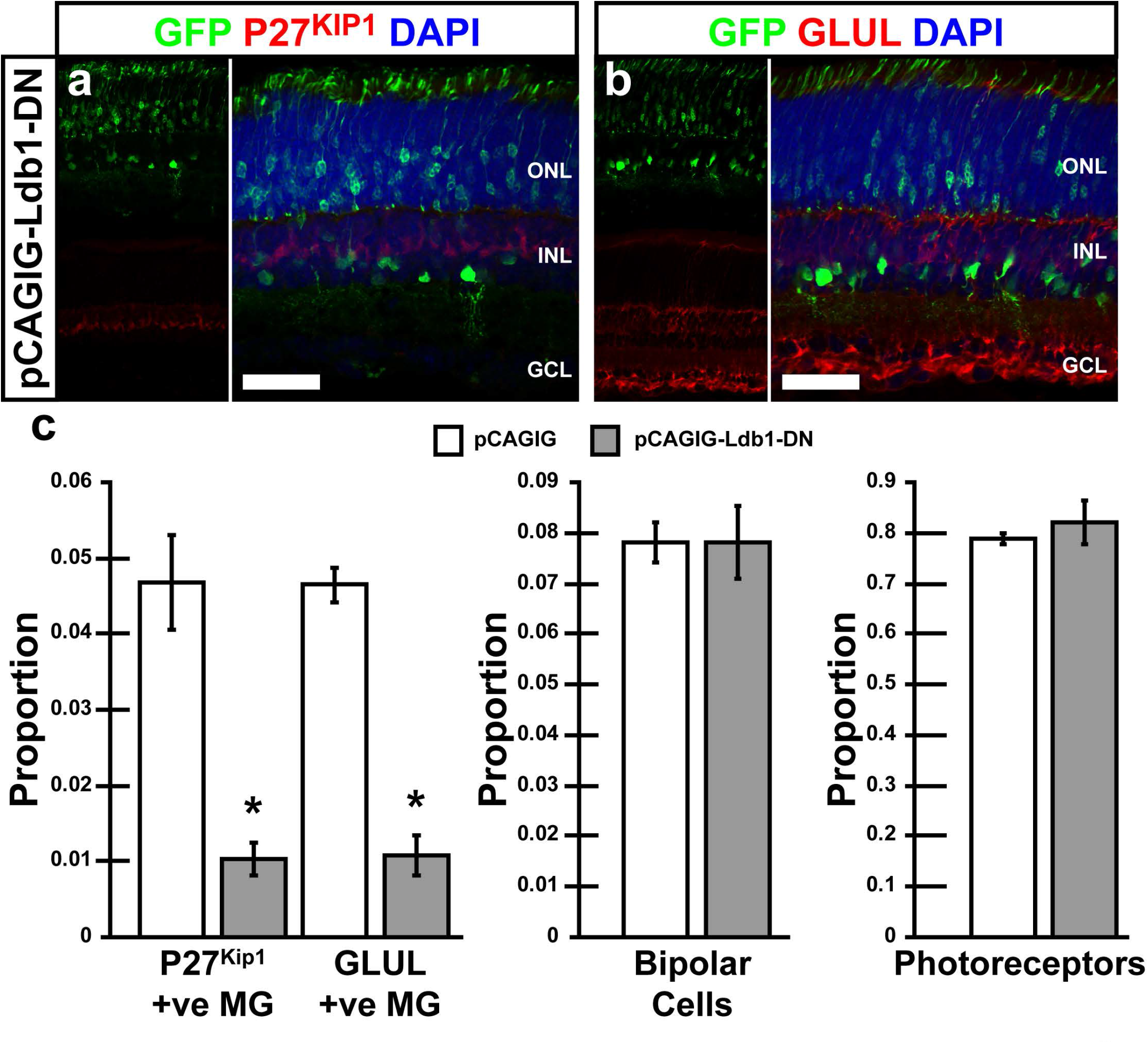
Electroporation of a dominant-negative *Ldb1* construct (pCAGIG-Ldb-DN) phenocopies *Lhx2* loss of function in postnatal retina. (a, b) Electroporation of pCAGIG-Ldb1-DN at P0 by electroporation resulted in a significant decrease at P14 of MG (P27^Kip1^ and GLUL +ve). (c) Quantification of MG (P27^Kip1^ and GLUL +ve), bipolar cells, and photoreceptors in pCAGIG vs. pCAGIG-Ldb1-DN electroporated retinas. Scale bars, 50 ¼m (a, b).

**Supplemental Figure 7.**
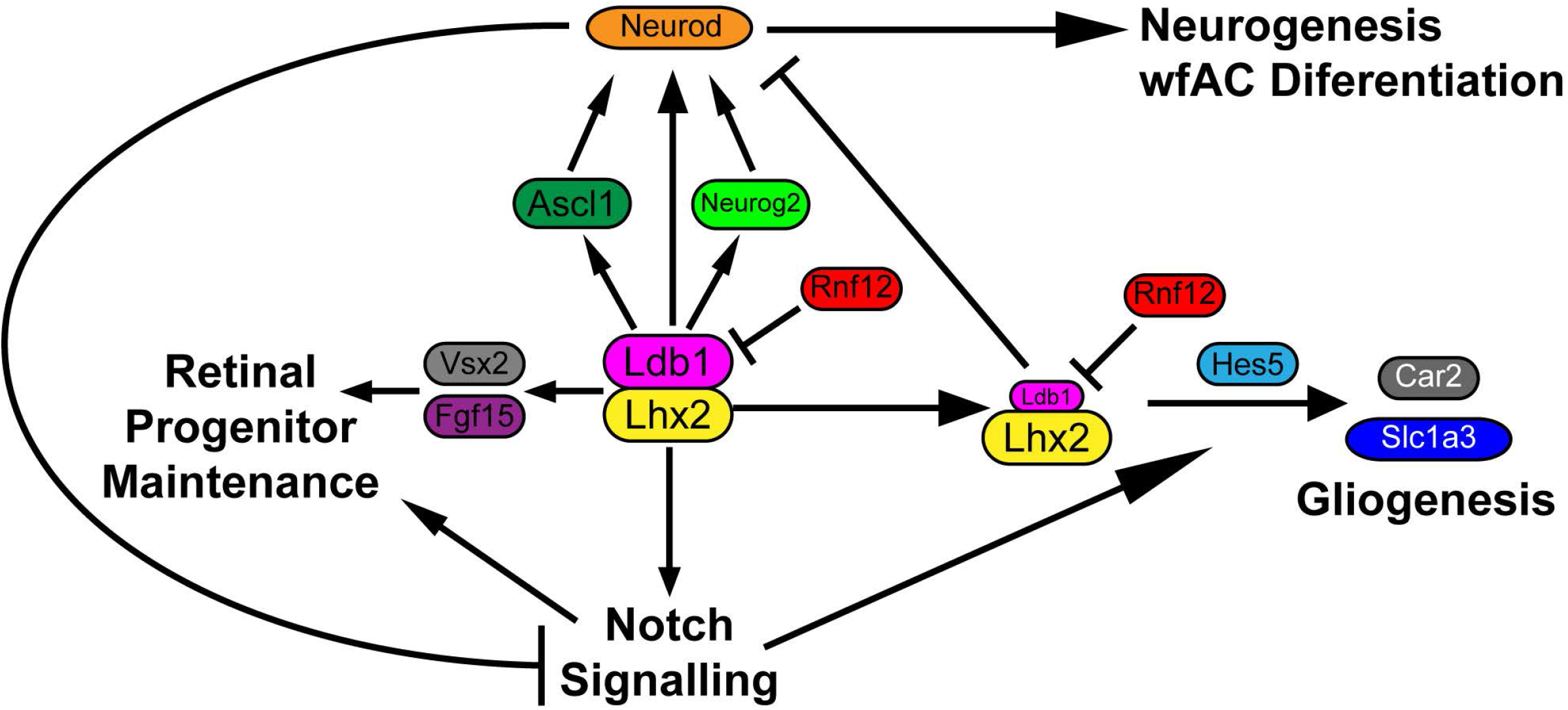
Model of Lhx2-dependent regulation of retinal gliogenesis. Lhx2, in combination with Ldb1, directly activates expression of genes that promote Notch signaling, proliferation, and both neurogenic and gliogenic competence in late-stage RPCs. The Lhx2/Ldb1 transcription activator complex also enhances expression of neurogenic bHLH factors, which in turn feed back to inhibit Notch signaling and drive neurogenesis. Rnf12, which selectively degrades Ldb1 via its E3 ubiquitin ligase activity, inhibits Lhx2/Ldb1-dependent activation of neurogenic differentiation, facilitating a transition to Müller gliogenesis.

**Supplemental Table 1:**

RNA-Seq data from P0.5 *Pdgfra-Cre;Lhx2^lox/lox^* retina was previously described in (1). RPKM values for each gene are listed.

